# Offspring chemical control of adult reproductive transitions in a social insect

**DOI:** 10.1101/2025.10.11.680753

**Authors:** Baptiste Piqueret, Jerrit Weissflog, Sandra Tretter, Tim Zetzsche, Daniel Veit, Stefan Bartram, Rayko Halitschke, Yuko Ulrich

## Abstract

Parental care enhances offspring survival and growth but often entails a trade-off in which caregivers temporarily suppress their own reproduction to invest in existing young. In vertebrates, these parental reproductive cycles are controlled by offspring-derived cues that reduce or suppress parental fertility. While many insects display obligate parental care, the role of offspring cues in regulating adult reproduction remains unresolved outside advanced eusocial taxa, where reproductive cycles have largely been lost. Here, we investigate reproductive cycles in the clonal raider ant *Ooceraea biroi*, in which totipotent females alternate between caring for larvae and laying eggs. Using custom behavioral assays, we show that larvae inhibit adult reproduction without physical contact, implicating volatile cues. Chemical analyses identified a previously undescribed larva-specific compound, methyl 3-ethyl-2-hydroxy-4-methylpentanoate (MEHMP), which is absent from other developmental stages and ant species. Exposure to synthetic MEHMP recapitulated the inhibitory effect of larval volatiles, confirming its role as a pheromone that suppresses adult egg laying. To our knowledge, this is the first brood pheromone described in ants. By providing a direct chemical link between offspring presence and parental reproductive suppression, our findings underscore the central role of offspring signals in mediating parental reproductive investment across animals that care for their young.

**Significance Statement:** Parental care enhances offspring survival but induces profound physiological changes in caregivers. A common feature across animals is that adults temporarily suppress their own reproduction while caring for the young. The offspring cues enforcing these cycles remain poorly understood, especially in insects. Using the clonal raider ant *Ooceraea biroi*, in which all females alternate between reproduction and brood care, we identify a larval pheromone that suppresses adult egg laying. This pheromone is a previously undescribed volatile chemical compound, and exposure to its synthetic version mimics the full biological effect of larvae volatiles. By providing a direct link between offspring presence and adult reproductive suppression, our work highlights how offspring can chemically control the reproduction of their caregivers.

## Introduction

Parental care provides significant benefits by increasing offspring survival and growth, while eliciting profound physiological and behavioral changes in caregivers. In species with offspring that require extended care, parental care often induces a trade-off wherein caregivers temporarily suppress their reproductive activity. This phenomenon occurs across diverse taxa, suggesting an advantage to prioritizing existing offspring over new reproductive attempts.

In many vertebrates, the behavioral sequence that separates reproduction and brood care is controlled by offspring-derived cues. For example, tactile cues associated with suckling contribute to lactational infertility in mammals (1, 2). Similarly, tactile cues associated with egg incubation reduce egg laying in many birds (3–5). The temporary suppression of reproduction is lifted when these offspring-derived cues decrease or disappear, e.g., at weaning, allowing parents to resume reproductive activity.

Although many insects exhibit obligate parental care (6), the role of brood-derived cues in regulating reproductive transitions in adult caregivers remains largely unresolved (7, 8). Current knowledge on insect brood-derived cues is largely based on advanced eusocial insects, in which queens monopolize reproduction and brood care is restricted to workers. In these systems, individual reproductive transitions no longer occur because egg laying and caregiving are performed by separate, morphologically specialized colony members, who have largely lost the ability to switch between reproductive and caregiving roles. Accordingly, the only brood-derived cues known to affect adult reproduction, the honeybee brood pheromones (*E*)-β-ocimene (9) and a mix of ten esters (10), reinforce reproductive division of labor by suppressing worker ovarian development but do not regulate individual reproductive transitions. Consistent with the requirement for permanent reproductive suppression of workers, even brood stages that do not require active care (e.g., capped brood) produce brood pheromones (11, 12), and several of its active components are also produced by queens (13, 14), suggesting inbuilt redundancy in the enforcement of worker sterility. Thus, even in the absence of brood-derived cues, honeybee workers fail to restore full reproductive potential, underscoring the extent to which reproductive plasticity has been lost in advanced social taxa. To uncover brood cues regulating adult reproductive transitions, it is therefore advantageous to investigate systems that display more reproductive plasticity.

For this purpose, we used the clonal raider ant *Ooceraea biroi*. In this species, all females are totipotent and regularly alternate between reproductive and brood care states. This cycle is regulated by larvae, which inhibit reproduction and induce brood care behavior in adults (15–17). Because the presence of larvae in a colony is periodic, entire colonies alternate between brood care phases (larvae are present, workers do not lay eggs) and reproductive phases (larvae are absent, workers lay eggs). Furthermore, clonal raider ant colonies are naturally queenless, eliminating the confounding effects of queen pheromones, the primary mechanism of worker reproductive suppression in most social insects (18).

To investigate how larvae regulate adult reproductive transitions in *O. biroi*, we first developed a custom behavioral assay to ask whether physical contact and/or volatile cues mediate the inhibitory effect of larvae on adult egg laying. These assays strongly suggest a role for volatile chemical cues in controlling the switch between reproductive and brood care states in adults. Using chemical analysis of headspace volatiles of all brood stages, we identify the compound methyl 3-ethyl-2-hydroxy-4-methylpentanoate (MEHMP), which is exclusively produced by *O. biroi* larvae. Behavioral assays show that chemically synthesized MEHMP inhibits adult egg laying to the same extent as larval volatiles. Collectively, these studies implicate a larval pheromone in the regulation of reproductive transitions.

## Results

We designed a behavioral platform to quantitatively study the influence of brood on adult egg laying. This platform utilized multiplexed 2-chamber arenas with controlled airflow, thereby allowing the passage of volatile compounds but preventing physical contact of the ants between the chambers (Fig. 1A). Using this assay, we first asked whether larvae can inhibit adult egg laying without physical contact. To this aim, we placed live larvae in the first chamber and a small colony of 12 workers in the second chamber (“larvae distance” treatment). As controls, workers were placed in physical contact with larvae (“larvae contact” treatment) in the second chamber (mimicking the brood care phase) or in physical contact with pupae (“no larvae” treatment) in the second chamber (mimicking the reproductive phase). Clonal raider ant stock colonies lay no eggs in the brood care phase and lay approximately one egg per ant in the reproductive phase. As expected, ants in physical contact with larvae laid no eggs (median (interquartile range IQR): 0 (0–0) eggs per worker; Fig. 1B) and, in contrast, ants in physical contact with pupae laid eggs (1.08 (1.00–1.17) eggs per worker). Thus, our custom assay recapitulates the phenotype of stock colonies in the two phases of their reproductive cycle (Dunn’s test, larvae contact – no larvae: Z = -5.69, p < 0.001). Although the presence of larvae in the first chamber did not fully recapitulate the effect of contact with larvae (larvae contact – larvae distance: Z = -2.84, p < 0.01), this arrangement strongly reduced adult egg laying (0.46 (0.40–0.59) eggs per worker, ca. 58% reduction; larvae distance – no larvae: Z = -2.84, p < 0.01). This suggested that larvae reduce adult reproduction through volatile chemical compounds and/or that pupae stimulate adult egg-laying. We thus next investigated volatile compounds produced by larvae. To account for a potential pupal stimulation of adult egg-laying, pupae were included in all treatments in subsequent behavioral experiments.

**Figure 1:**
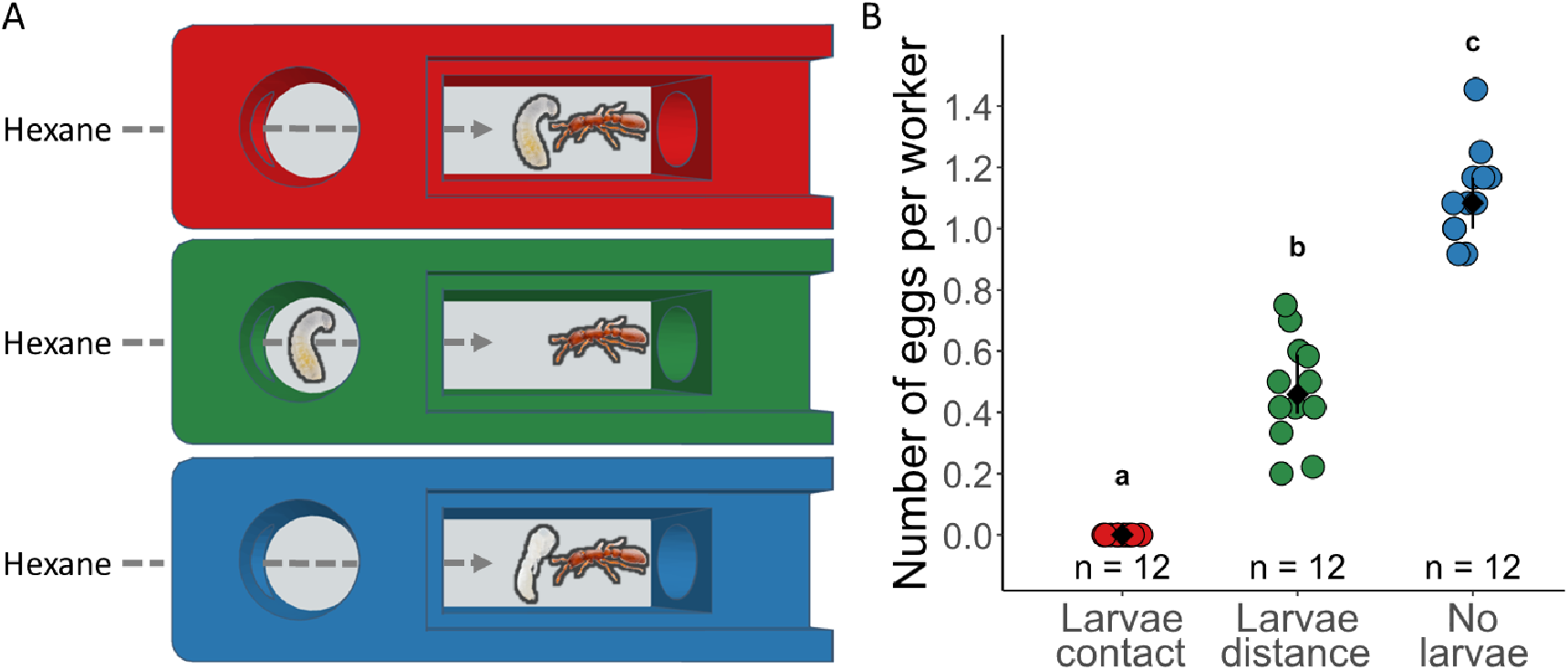
Larvae inhibit adult egg laying without direct contact. **(A)** 2-chamber arenas with controlled airflow. Treatments from top to bottom: workers in physical contact with larvae (red, “larvae contact”), workers without physical contact with larvae (green, “larvae distance”), workers in physical contact with pupae (blue, “no larvae”). All treatments were exposed to hexane to ensure comparability between experiments in this study. **(B)** Number of eggs per worker on the last experimental day. Data points represent replicate arenas. Black: median ± IQR. Different letters represent significantly different treatments (p < 0.05).

To identify candidate volatile compounds responsible for the observed effect on adults, we collected volatiles in the headspace of different *O. biroi* brood development stages and of other ant species using solid phase micro extraction (SPME) and analyzed them using gas-chromatography mass-spectrometry (GC-MS). We reasoned that any compound responsible for the observed effect should be present in headspace of larvae (the only brood stage inhibiting adult egg laying) and absent in the headspace of other brood stages (i.e., eggs, prepupae, pupae) of *O. biroi*. Because *O. biroi* feeds on other ant species (19), any candidate compound should also be absent in other ant species, so that *O. biroi* reproduction would not be inhibited by its prey. Using this approach, we identified a single larva-specific compound with a retention time of 8.8 min, which was produced only by *O. biroi* larvae (Fig. 2A), but not by alien ant brood (*Tetramorium bicarinatum* and *Iridomyrmex purpureus*, Fig. S1). The mass spectrum showed a strong *m/z* 90 base peak, which was used in extracted ion chromatograms for detailed comparisons and compound quantification (Fig. 2A, Figs. S1–S3). The *m/z* 90 ion is a diagnostic fragment of a McLafferty rearrangement of α-hydroxycarboxylic acid methyl esters, and initial library searches and further fragment analysis suggested a branched 8-carbon carboxylic acid methyl ester. We synthesized several isomers with different branching patterns and identified the larval volatile compound as methyl 3-ethyl-2-hydroxy-4-methylpentanoate (MEHMP), a previously undescribed compound (Fig. 2B). The synthesized standard showed identical retention behavior as the natural compound on different GC stationary phases, as well as the same mass spectrum (Fig. 2C).

**Figure 2.**
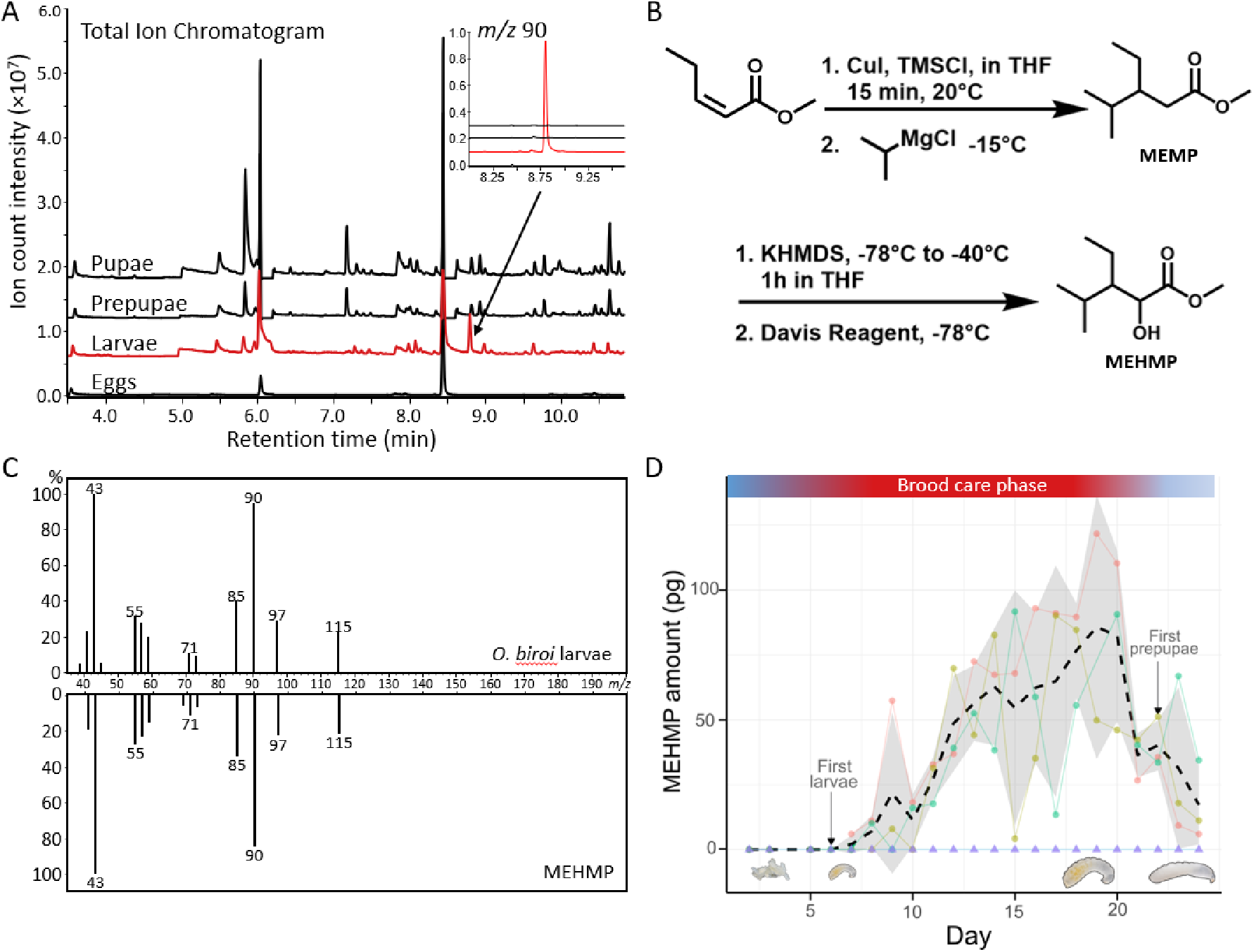
Identification of MEHMP as a larva-specific candidate pheromone. **(A)** GC-MS chromatograms of SPME headspace extracts of different *O. biroi* brood stages (red: larvae). Top insert: 90 *m/z* extracted ion chromatograms. **(B)** Synthesis of MEHMP from methyl (*Z*)-pent-2-enoate. The reaction conditions for the subsequent 1,4-Michael-addition (using copper(I)-iodide and isopropyl magnesium chloride) and α-hydroxylation (using a Davis-oxidation) are shown and produced the target compound in 55% yield over the two steps. **(C)** Validation of the identification using MS spectra of the volatile produced by larvae (top) and synthetic MEHMP (bottom). **(D)** Daily abundance of MEHMP tracks the *O. biroi* colony cycle. Circles: replicate colonies of 100 workers in plaster nests (n = 3); triangles: control, empty plaster nests (n = 2). Dashed line: mean of ant colonies; grey: s.d. Top bar: transitions between reproductive phases (blue) and brood care phase (red).

If MEHMP controls individual and colony phasic reproduction, we would expect its production to correlate with natural brood development and colony phases. To test this, we quantified daily MEHMP abundances in the headspace of colonies of 100 ants that naturally progressed from the reproductive to the brood care phase, and back to the reproductive phase, using volatile collection on polydimethyl siloxane (PDMS) tubing (Fig. 2D). We found that the abundance of MEHMP closely mirrored cycle phases and brood development. It was undetectable in colonies as adults laid eggs, appeared after eggs hatched into larvae, increased in abundance as larvae grew over the course of the brood care phase, and waned shortly before larvae became prepupae, which marks the cessation of larval feeding and the start of a new reproductive phase. Thus, life stage specific (Fig. 2A) and colony-level, temporal (Fig. 2D) measurements show that the production of MEHMP is specific to larvae and tracks colony cycles, making it a viable candidate larval pheromone controlling adult reproductive transitions and colony reproductive cycles.

The amount of MEHMP collected in the larval headspace was quantified using calibration curves generated from PDMS (Fig. S2) and SPME (Fig. S3) sampling of dilution series of synthetic MEHMP. This analysis indicated that a 4^th^ instar *O. biroi* larva produces approximately 1–2 pg of MEHMP per day (PDMS, day 19: mean ± s.d.:1.07 ± 0.63 pg day^-1^, n = 2; SPME: 2.13 ± 0.12 pg day^-1^, n = 3; see Methods). The synthesized MEHMP is a racemic mixture composed of four stereoisomers (Fig. S4). While the absolute configuration of each isomer could not be resolved at that time, GC-MS analyses with a chiral stationary phase suggest a stereospecific production of MEHMP by the larvae. Larval production of MEHMP is dominated (>90%) by one stereoisomer. This isomer accounts for only 34% of the synthetic mixture.

We next used our behavioral platform to assess whether MEHMP affects adult reproduction (Fig. 3). Ant groups were exposed to 200 pg of synthetic MEHMP per day. Due to the racemic mixture of the synthesized MEHMP, this quantity represents ca. 68 pg of the main stereoisomer produced by larvae, which is comparable to ca. 35–71 larvae equivalents (Fig. S5, “MEHMP”). The MEHMP-exposed ant groups were compared to the same groups used previously, i.e. ants in physical contact with larvae (“larvae contact”), ants without physical contact with larvae (“larvae distance”), and ants in physical contact with pupae only (“no larvae”, Fig. 3A). As described above, pupae were included in all the treatments to control for potential pupal stimulation of egg laying. Exposure to MEHMP alone induced a ca. 33% reduction in egg laying compared to control (MEHMP: median (IQR): 0.67 (0.54–0.75) eggs per worker, no larvae: 1.00 (0.92–1.04) eggs per worker; MEHMP – no larvae: Z = -4.39, p < 0.001; Fig. 3B) and recapitulated the effect of exposure to the full blend of larval volatiles (larvae distance: 0.67 (0.50–0.77) eggs per worker; MEHMP – larvae distance: Z = -0.84, p = 0.40; larvae distance – no larvae: Z = -3.63, p < 0.001). Latency to lay eggs did not vary across treatments in which eggs were laid (Kruskal-Wallis test: *χ*^2^ = 4.13, d.f. = 3, p = 0.25), indicating that egg laying was in fact inhibited and not merely delayed.

**Figure 3:**
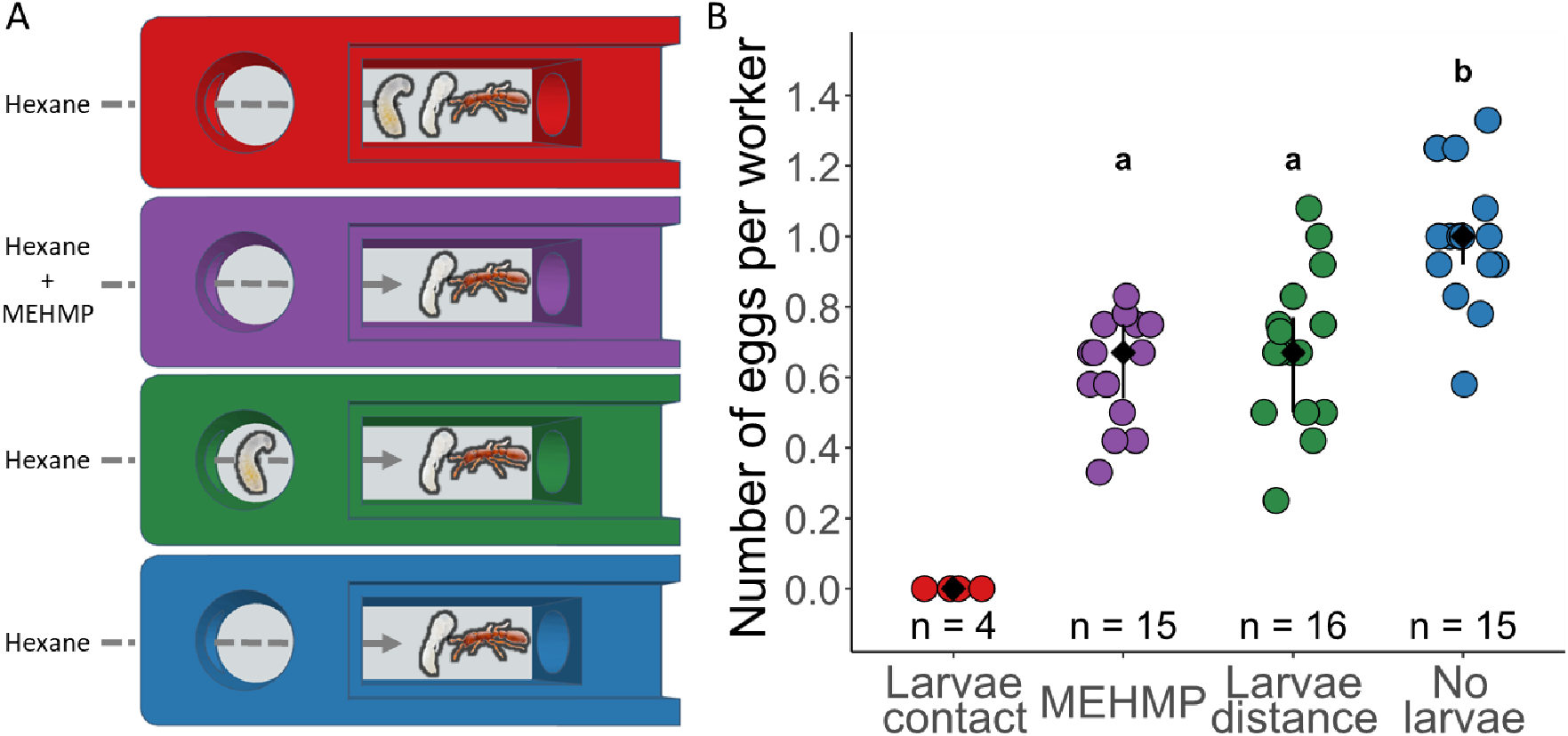
MEHMP inhibits adult egg laying. **(A)** 2-chamber arenas with controlled airflow. Treatments from top to bottom: workers in physical contact with larvae (red, “larvae contact”), workers exposed to MEHMP (purple), workers without physical contact with larvae (green, “larvae distance”), workers not exposed to larvae (blue, “no larvae”). Ants in all treatments were exposed to hexane and were in direct contact with pupae to ensure comparability among treatments. **(B)** Number of eggs per worker on the last experimental day. Data points represent replicate arenas. Black: median ± IQR. Different letters represent significantly different treatments (p < 0.05).

Additionally, to test the specificity of the effect of MEHMP, we included a treatment with 200 pg per day of its non-hydroxylated synthetic precursor, methyl 3-ethyl-4-methylpentanoate (MEMP), which is absent from the headspace of *O. biroi* larvae but was previously identified as a pheromone in another ant (20). Exposure to MEHMP inhibited adult egg laying more strongly than MEMP, indicating that the response is compound-specific and the hydroxylation modification plays a role in the pheromone function (Fig. S5; MEMP: 0.83 (0.67–0.94) eggs per worker, MEHMP – MEMP: Z = -2.38, p < 0.05).

## Discussion

Here, we show that a previously undescribed volatile compound, produced exclusively by larvae, regulates reproductive transitions in adult *O. biroi* ants by inhibiting egg laying. Because larvae occur only periodically in colonies, this pheromone likely contributes to reproductive synchrony, coordinating colony-level transitions between brood care and reproductive phases. To our knowledge, this represents the first identified larval pheromone that regulates reproductive cycles in an insect, and the first brood pheromone reported in ants, whose existence has until now remained uncertain (21, 22). While MEHMP was previously undescribed, the structurally related MEMP was identified in adult *Formica* and *Polyergus* ants (20, 23), though this compound likely has a different function in these species. In our experiment, exposure to MEMP did not recapitulate the effects of MEHMP, showing the effects on adult reproduction are specific to the larval pheromone.

Past work in *O. biroi* has shown that larvae control adult ovarian activation by influencing adult insulin signaling (24). The presence of larvae inhibits adult reproduction by suppressing *insulin-like peptide 2* (*ilp2*). Experimentally elevating ILP2 in adults overrides larval suppression of worker reproduction, breaking the colony’s reproductive cycle. The larval pheromone (MEHMP) we have identified is a strong candidate as the proximate signal modulating ILP2 levels, functioning upstream of insulin-controlled endocrine regulation. In this light, decreasing adult sensitivity to this pheromone would likely disrupt the colony cycle, similar to the effect of artificially elevating ILP2. This integration of pheromone signaling with insulin-mediated reproductive control brings us closer to a complete mechanistic understanding of how a complex social interaction, larval control of adult reproduction, is orchestrated in this system.

The volatility of MEHMP likely represents an important requirement for ensuring colony-wide enforcement of reproductive synchrony. Like other social insects, *O. biroi* exhibits division of labor (25), with some workers (e.g., foragers) having limited direct contact with larvae. If reproductive inhibition relied solely on contact-based cues, such individuals could escape suppression, potentially disrupting colony-wide reproductive synchrony. A volatile signal ensures that all workers, regardless of their tasks, are exposed to larva-derived inhibition and remain in the non-reproductive state during the brood care phase. In this way, the pheromone provides a possible mechanistic basis for how *O. biroi* colonies maintain coordinated shifts between reproduction and brood care.

Both the larval pheromone of *O. biroi* and the brood pheromones of the honeybee inhibit adult reproduction, but they differ in the brood stages that produce them and in the colony members they affect. Honeybee brood pheromones are produced by all brood stages (11, 12) and act redundantly with queen pheromones to maintain continuous suppression of worker reproduction (9, 10). However, these pheromones do not suppress queen egg-laying and may even promote it (26). This enables brood care by workers and egg laying by queens to occur simultaneously. In *O. biroi*, by contrast, the pheromone is only produced by larvae, the brood stage that requires the most care (other brood stages do not feed). As a result, the pheromone transiently suppresses worker reproduction only when larvae are present, causing brood care and egg laying to occur sequentially rather than simultaneously. *O. biroi* therefore illustrates how brood cues can orchestrate coordinated, colony-wide reproductive transitions in a queenless social species in which all females are totipotent.

Although our study identifies a volatile larval pheromone as a central regulator of reproductive transitions in *O. biroi*, several questions remain. While the synthetic pheromone fully recapitulates the biological effect of larval volatiles, it only partially reproduces the effect of physical contact with larvae, suggesting additional contributions from other larval cues. Partial inhibition of worker reproduction is characteristic of chemically identified honeybee brood pheromones, where individual components of the brood ester pheromone and the volatile E-β-ocimene reduce ovary development by approximately 33% relative to controls (9, 10). The ca. 33% reduction in egg laying induced by MEHMP is therefore well within the range of established brood pheromone effects in social insects. Future work should determine whether other non-volatiles larval chemical cues or non-olfactory cues (e.g., tactile cues) contribute to adult reproductive suppression. At the mechanistic level, elucidating the sensory pathways by which the pheromone is perceived and how this perception is coupled to endocrine regulation of reproduction will be crucial for establishing a complete model. Finally, comparative studies in other ants, as well as subsocial and primitively eusocial species, could reveal whether larval pheromones represent an ancestral regulator of adult reproductive transitions or a derived innovation of *O. biroi*. Larvae suppress worker reproduction in at least one primitively eusocial bee (27) and three ant species (28–30), and they stimulate queen reproduction in at least two ant species (31, 32), making these promising systems in which to discover additional larval pheromones and compare their effects across castes and social structures.

In summary, we identify a larval pheromone that regulates adult reproductive transitions in an insect, providing a direct link between brood care and reproductive suppression. Our discovery adds ants to the growing list of taxa in which offspring-derived cues can suppress caregiver reproduction. Whereas offspring reduce parental fertility via tactile cues in many vertebrates, in *O. biroi* this occurs at least in part through chemical communication. Our findings underscore the central role of offspring cues in regulating adult reproduction across animals that care for their young.

## Materials and Methods

### Egg laying behavioral assay

We developed a platform to quantify the effects of brood, brood volatile organic compounds (VOCs), and synthetic compounds on adult egg laying behavior. The platform consisted of multiplexed 2-chamber arenas (65 × 20 mm) coupled with an odorant delivery system with controlled airflow. Arenas were 3D-printed in polylactic acid and comprised an upstream round chamber (diameter: 12 mm) closed with an airtight metallic lid and a downstream rectangular chamber (28 × 11 mm) closed with a clear acrylic lid, allowing egg counting without colony disturbance. The chambers were separated by a round metallic gate (diameter: 8.5 mm, width: 2 mm) that allowed air flow but prevented physical contact (including antennal contact) between ants in the different chambers. Both chambers had a humidified plaster-of-Paris floor. The input and output air flows of each arena were set by analogous flow controllers (Key Instruments, USA). Air passed through two 250 mL bottles filled with distilled water and one 250 mL odorant delivery bottle (see below) before entering the upstream chamber and carrying any volatile compounds from that chamber to the downstream chamber.

Experiment 1 (Fig. 1A). We tested whether larvae can inhibit adult egg laying without physical contact using three treatments: *i*) “larvae contact”: 12 adults (16-day old, genetic lineage A (33)) were placed in direct contact with 12 larvae (6-day old, genetic lineage B) in the downstream chamber; this treatment is analogous to the brood care phase, *ii*) “no larvae”: 12 adults were placed in direct contact with 12 pupae (5-day old, genetic lineage D) in the downstream chamber; this treatment is analogous to the reproductive phase, *iii*) “larvae distance”: 36 larvae were placed in the upstream chamber, and 12 adults in the downstream chamber. Twelve replicate arenas were used for each treatment. Batches of four arenas were connected to the same flow controller. The natural larvae-to-worker ratio in *O. biroi* colonies is ca. 1:1; in *(iii)*, we used a 3:1 larvae-to-worker ratio to increase the likelihood of capturing an effect, if it existed. This ratio was adjusted to the natural level in Experiment 2 (see below). Air was pushed into each arena at a flow of 75 mL min^-1^. For all arenas, a 4 mL vial of hexane solvent was placed in the odorant delivery bottle and exchanged daily. Hexane exposure was included to ensure it did not prevent ant survival and egg laying, with future odorant-delivery experiments in mind.

Experiment 2 (Fig. 3A). We tested whether the putative larval pheromone inhibits adult egg laying, using a treatment with methyl 3-ethyl-2-hydroxy-4-methylpentanoate (MEHMP) delivery in addition to the three treatments used in Experiment 1. To test the specificity of effects, we included a treatment with methyl 3-ethyl-4-methylpentanoate (MEMP) delivery, a synthetic precursor of MEHMP that is not produced by *O. biroi* larvae. As before, a 4 mL vial of hexane was placed in the odorant delivery bottle of each arena and exchanged daily. In contrast to Experiment 1, pupae were included in all groups to control for potential pupal stimulation of egg laying. Additionally, we introduced three adults in the upstream chamber of all treatments to act as caregivers for the larvae. The five treatments were: *i*) “larvae contact”: 12 adults (11-day old, genetic lineage B) were placed in direct contact with 12 larvae (6-day old, genetic lineage B) and 12 pupae (0-day old, genetic lineage B) in the downstream chamber; *ii*) “no larvae”: 12 adults were placed in direct contact with 12 pupae in the downstream chamber; *iii*) “larvae distance”: 12 larvae were placed in the upstream chamber, and 12 adults were placed in the downstream chamber with 12 pupae; *iv*) “MEHMP”: 12 adults were placed in the downstream chamber with 12 pupae; 200 pg of methyl 3-ethyl-2-hydroxy-4-methylpentanoate were added to the hexane vial, *v*) “MEMP”: 12 adults were placed in the downstream chamber with 12 pupae; 200 pg of methyl 3-ethyl-4-methylpentanoate were added to the hexane vial. Sixteen replicate arenas were used per treatment, except for (*i*), which had 4 replicates. Because ants in this treatment invariably laid no eggs in Experiment 1, we included this treatment as a positive control but excluded it from statistical analyses in Experiment 2, to allow sufficient replication in the four other treatments. Air was pushed into each arena at a flow of 75 mL min^-1^. Additionally, to minimize leaks of the synthetic compounds, air was actively pumped out of each arena at 87.5 mL min^-1^ and filtered through activated carbon before release into the experimental room.

In both experiments, the survival of adults, larvae, and pupae, as well as the number of eggs laid was monitored daily. Dead larvae and pupae were replaced each day to maintain constant brood exposure; dead adults were removed but not replaced. Arenas were humidified daily but received no food to ensure that any differences in egg laying were not driven by differences in food intake or foraging behavior. We calculated the median latency to lay eggs in the negative control (“no larvae”) and ended the experiments 3 days later, which occurred on day 8 in Experiment 1 and on day 9 in Experiment 2.

In both experiments, adult survival was high (>93% in Experiment 1, >95% in Experiment 2). Arenas in which >33% of adults died or escaped were excluded from further analyses, resulting in the exclusion of no arenas in Experiment 1 and two arenas in Experiment 2.

### Identification of the candidate pheromone

To identify candidate volatile compounds responsible for adult reproductive inhibition, we collected volatiles in the headspace of different life stages using solid phase micro extraction (SPME) and analyzed them using gas chromatography-mass spectrometry (GC-MS). We incubated 50–100 larvae (n = 18, mixed instars), 200 eggs (n = 3), 50–200 prepupae (n = 5) and 100 pupae (n = 7) in 1.5 mL glass vials with a SPME fiber (DVB/CAR/PDMS, Supelco, USA) inserted through the septum cap (PTFE, Macherey-Nagel, Germany) and exposed them to the brood headspace for 18 h at 28 °C. Additionally, we used the same method to collect headspace extracts from the brood of two other ant species, *T. bicarinatum* (15 larvae, n = 1; 10 pupae, n = 3) and *I. purpureus* (3–10 larvae, n = 5; 2 pupae, n = 1), maintained at 25 °C in a separate room. After sampling, the fiber was desorbed by splitless injection at 230 °C into a Shimadzu QP2010 GC-MS, equipped with a SLB-5ms GC column (30 m × 0.25 mm × 0.25 μm, Supelco, USA) and helium as carrier gas (1.2 mL min^−1^). The oven temperature was programmed at 50 °C for 1 min, then increased to 260 °C at 10 °C min^−1^, then to 320 °C at 20 °C min^−1^ and held for 5 min. Mass spectra were recorded with electron impact ionization at 70 eV. The software GCMSsolution (v4.20, Shimadzu Corporation) was used for data analysis. Identification of the compounds was accomplished by comparison of library databases, diagnostic ions and retention indices (RI) calculated with a standard n-alkane solution (C7–C30, 99%, Sigma-Aldrich). The candidate pheromone shows a RI of 1152 on the nonpolar SLB-5ms phase, and additional analyses on a polar phase (ZB-WAX, Phenomenex) yielded a RI of 1635. To separate stereoisomers, additional SPME samples were analyzed by GC-MS on a chiral GC column (CYCLOSIL-B, 30 m × 0.25 mm × 0.25 μm, Agilent Technologies, USA) with a flow rate of 1.2 mL min^-1^ and an oven temperature gradient of 40 °C for 1 min, increased to 150 °C with 5 °C min^-1^, then with 20 °C min^-1^ to 250 °C and a final hold at 250 °C for 2 min.

### Chemical synthesis

Commercially available chemicals were used without further purification. All reactions were carried out under an argon atmosphere. Preparative column chromatography was performed on Silica gel 60 (230–400 mesh, Carl Roth GmbH) and TLC analysis on commercial Merck silica gel 60 F_254_ plates. NMR spectra were measured on a Bruker Avance 400 NMR spectrometer. Chemical shifts are reported in ppm downfield from TMS.

#### Synthesis of methyl 3-ethyl-4-methylpentanoate (MEMP)

A 250 mL three-necked flask, equipped with rubber septum and dropping funnel, was charged with 1.18 g copper(I)-iodide (6.1 mmol) and put under an atmosphere of argon. The flask was loaded with 70 mL anhydrous THF and 7 g of methyl (*Z*)-pent-2-enoate (61 mmol) were added. Chlorotrimethylsilane (9.4 mL, 74 mmol) was then added via syringe. After stirring for 15 min at 20 °C, the mixture was cooled to -15 °C in an ice/methanol bath. A 2 M solution of isopropylmagnesium chloride in THF (36 mL, 74 mmol, Merck) was added dropwise over 2 hours, while keeping the temperature below -10 °C. After stirring for an additional hour at -15 °C, the reaction was slowly quenched by adding methanol (5 mL) and 2 M ammonium chloride solution (100 mL). After separation of the liquid layers, the aqueous phase was extracted twice with ∼75 mL diethyl ether. The organic phases were pooled and washed with water and brine (100 mL each), dried with anhydrous MgSO_4_, filtered and concentrated in vacuo. Vacuum distillation gave MEMP (7.5 g, 76% yield) as a colorless liquid (B.p.110–120 °C at 50 mbar). ^1^H-NMR: (400 MHz, CDCl_3_) d = 3.51 (s, 3H), 2.07 (qd, *J*_*1*_ = 21.9 Hz, *J*_*2*_ = 6.7 Hz, 2H), 1.50-1.60 (m, 2H), 1.01-1.20 (m, 2H), 0.89 (m, 9H) ppm; ^13^C-NMR: (100 MHz, CDCl_3_) d = 174.7, 51.4, 42.6, 35.6, 29.4, 23.7, 19.4, 18.5, 11.6 ppm. EI-MS: *m/z* (rel. int.) = 127 (10), 115 (13), 101 (6), 87 (31), 85 (33), 83 (18), 74 (100), 70 (23), 59 (18), 57 (16), 43 (27).

#### Synthesis of methyl 3-ethyl-2-hydroxy-4-methylpentanoate (MEHMP)

MEMP (460 mg, 2.9 mmol) was dissolved in 6 mL anhydrous THF. The solution was cooled to -78 °C and a 1 M solution of potassium bis(trimethylsilyl)amide (4 mL, 40 mmol) was added via syringe. The mixture was allowed to warm to -40 °C and stirred for 1 h at this temperature before being cooled to -78 °C again. Davis oxaziridine (3-phenyl-2-phenylsulfonyl-1,2-oxaziridine, 900 mg, 3.5 mmol) was dissolved in 2 mL anhydrous THF and added via syringe to the reaction mixture. After stirring for 1 h at -78 °C, the mixture was quenched with 2 M ammonium chloride solution (10 mL).

The mixture was vigorously stirred at room temperature for 1 h, before separation of the liquid layers. The aqueous phase was extracted with diethyl ether (2 × 10 mL), and the combined organic phases were washed with water and brine (10 mL each), before being dried with anhydrous MgSO_4_, filtered and concentrated in vacuo. Column chromatography (silica gel, 9:1 n-hexane/ethyl acetate v/v as eluent) gave MEHMP (370 mg, 73% yield) as a colorless oil. GC-MS-analyses (WAX-column) and NMR analysis showed that two pairs of diastereomers are present in a ratio of 8:3.

Diastereomer A: ^1^H-NMR: (400 MHz, CDCl_3_): d = 4.34 (dd, *J*_*1*_ = 4.9 Hz, *J*_*2*_ = 2.2 Hz, 1H), 3.786 (s, 3H), 2.72 (d, *J* = 4.96 Hz, 1H, (OH)), 1.80-1.90 (m, 1H), 1.30-1.60 (m, 3H), 0.98 (d, *J* = 6.8 Hz, 6H), 0.86 (t, 3H) ppm; ^13^C-NMR: (100 MHz, CDCl_3_) d = 176.9, 71.2, 52.4, 49.3, 29.2, 20.6, 20.0, 19.4, 12.8 ppm. EI-MS: *m/z* (rel. int.) = 115 (23), 97 (26), 90 (100), 85 (44), 73 (9), 71 (14), 59 (24), 57 (31), 55 (30), 43 (86), 41 (30).

Diastereomer B: ^1^H-NMR: (400 MHz, CDCl_3_): d = 4.30 (dd, *J*_*1*_ = 4.7 Hz, *J*_*2*_ = 3.8 Hz, 1H), 3.794 (s, 3H), 2.65 (d, *J* = 5.15 Hz, 1H, (OH)), 1.68-1.78 (m, 1H), 1.30-1.60 (m, 3H), 0.96 (t, *J* = 6.5 Hz, 3H), 0.91 (d, *J* = 6.5 Hz, 3H), 0.90 (t, 3H) ppm; ^13^C-NMR: (100 MHz, CDCl_3_) d = 176.7, 72.0, 52.37, 49.2, 28.4, 21.9, 19.38, 19.2, 12.5 ppm. EI-MS: *m/z* (rel. int.) = 115 (23), 97 (26), 90 (100), 85 (44), 73 (9), 71 (14), 59 (24), 57 (31), 55 (30), 43 (86), 41 (30).

### Time-course experiment and candidate compound quantification

Three groups of 100 adult ants (mixed age, genetic lineage B) were placed in circular, airtight plastic cups (height: 4 cm, base diameter: 3 cm, top diameter: 4 cm) with a humidified plaster-of-Paris floor and kept at 28 °C. Once ants began laying eggs, nest headspace was collected daily using 5-mm pieces of polydimethyl siloxane (PDMS) tubing suspended in the nest on a pin for 24 h. As a control, headspace was collected daily from identical plaster-lined cups without ants (n = 2). When eggs hatched into larvae, marking the start of the brood care phase, each colony was fed frozen brood of *T. bicarinatum* every 3–4 days. The experiment ended 2 days after the appearance of the first prepupae. We estimated that ca. 70–80 larvae were present in each nest during the experiment. PDMS headspace samples were inserted into a Thermal Desorption (TD) unit sampling tube and analyzed on a Shimadzu QP2010 GC-MS equipped with a TD-20 thermal desorption system and an SLB-5ms GC column (30 m × 0.25 mm × 0.25 μm; Supelco, USA). The carrier gas was helium (1 mL min^−1^). Injections were splitless (230 °C). The oven program was: 50 °C for 1 min, ramp to 260 °C at 10 °C min^−1^, ramp to 320 °C at 20 °C min^−1^ and hold at 320 °C for 5 min. Mass spectra were recorded with electron-impact ionization at 70 eV.

The amount of MEHMP collected in the headspace was quantified using calibration curves. 1 µL of a hexane solution containing 0 pg, 50 pg, 100 pg, 500 pg, or 1 ng of MEHMP (n = 3 for each concentration) was added to the bottom of a 1.5 mL glass vial, with 5-mm pieces of PDMS tubing suspended on a pin through the cap for 18 h. Samples were analyzed as above. A calibration curve was generated from the dilution series of the synthetic standard (peak area of the *m/z* 90 extracted ion chromatogram) and concentrations of the natural samples calculated (Fig. S2). This analysis indicated that each larva produced ca. 1 pg day^-1^ of MEHMP at the peak of production when larvae were 12–13 days old, on experimental day 19 (mean ± s.d.:1.07 ± 0.63 pg day^-1^, n = 2).

To confirm this estimate, we used SPME fibers sampled from 4^th^ instar larvae (n = 3, 100 larvae) (see “Identification of the candidate pheromone” above) and an independent calibration curve (Fig. S3). Here, 1 µL of a hexane solution containing 0 pg, 50 pg, 100 pg, or 250 pg of MEHMP (n = 3 for each concentration) was placed at the bottom of a 1.5 mL vial and exposed to a SPME fiber (DVB/CAR/PDMS) for 18 h. This analysis yielded an estimate of ca. 2 pg day^-1^ per larva (mean ± s.d.: 2.13 ± 0.12 pg day^-1^, n = 3), consistent with the PDMS-based quantification.

### Statistical analyses

All analyses were performed in R v4.5.1 (34). Kruskal-Wallis tests were used to compare egg laying across treatments on the last experimental day. To account for adult mortality, we modelled the number of eggs per worker (eggs per arena divided by the number of live adults). Pairwise comparisons between treatments were performed using Dunn’s tests (*dunn*.*test* function from package *dunn*.*test* (35)) with Benjamini-Hochberg adjustment for multiple comparisons. Kruskal-Wallis tests were used to compare latency to lay eggs across treatments.

## Acknowledgments

We thank Martin Kaltenpoth, Sarah O’Connor and Tobias Koellner for helpful discussions, Jonghyun Park for ant pictures, Sybille Lorenz for technical assistance with chemical analyses, Danny Kessler for access to climate chambers, and the Lise Meitner Research Group Social Behaviour for support with experiments. This is Clonal Raider Ant Project paper no. 40.

