## Supplementary Information - Fig.S1-S5 for "Offspring chemical control of adult reproductive transitions in a social insect"

**This PDF file includes:**

Figures S1 to S5

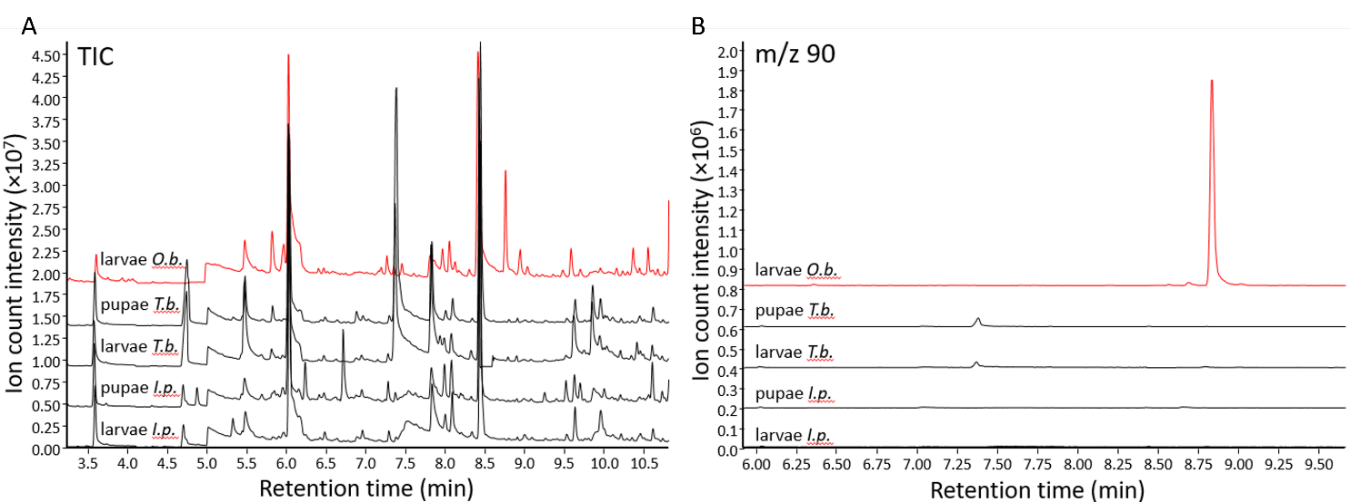

**Figure S1**. **(A)** GC-MS chromatograms of SPME headspace extracts of larvae and pupae of *Tetramorium bicarinatum* (*T.b.*, 15 larvae, 10 pupae) and *Iridomyrmex purpureus (I.p.*, 3 larvae, 2 pupae). The headspace profile of larvae from *O. biroi (O.b.)* is shown as a reference (red). **(B)** *m/z* 90 extracted ion chromatograms.

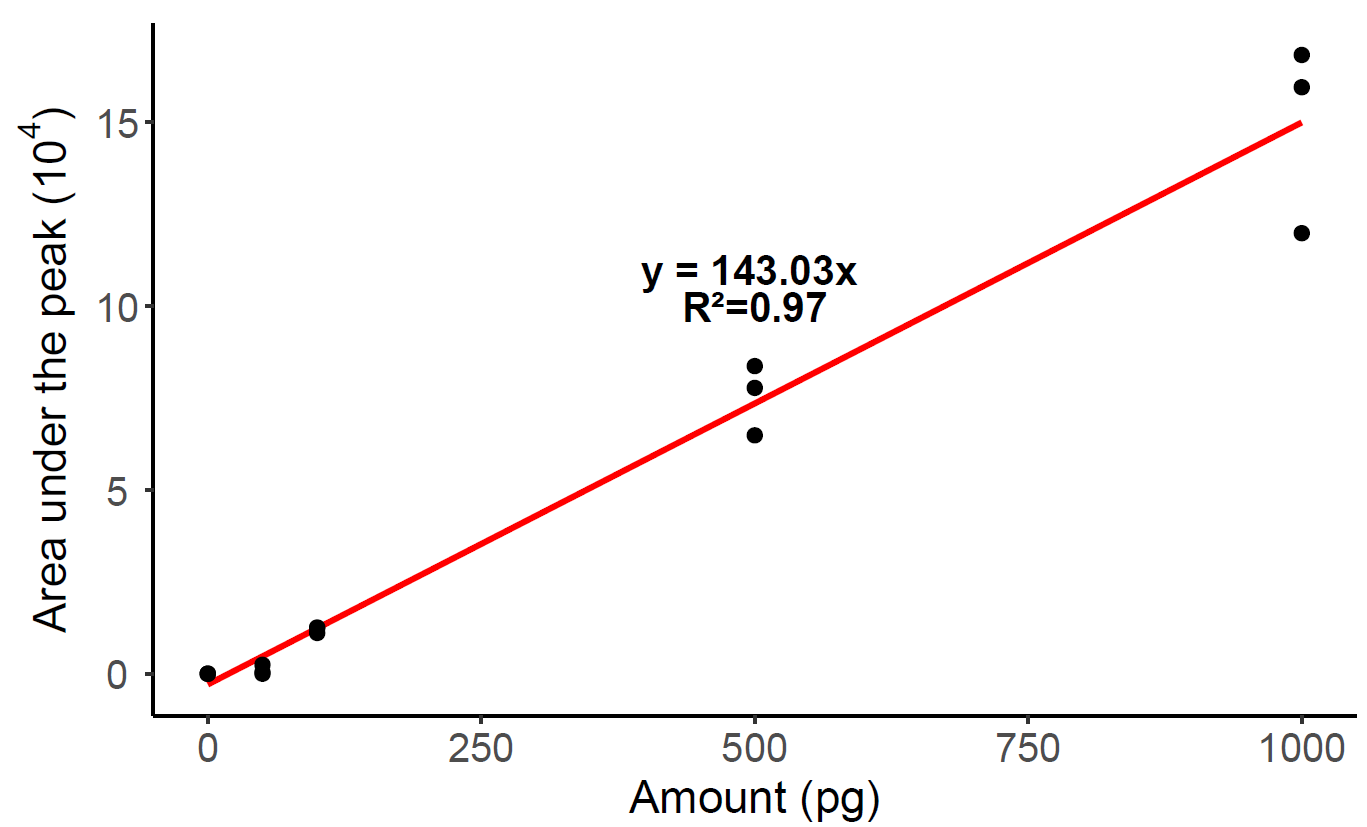

**Figure S2.** Calibration curve of methyl 3-ethyl-4-methyl-2-hydroxy-pentanoate (MEHMP) using PDMS and TD-GC-MS. 1 µL of a hexane solution containing 0 pg, 50 pg, 100 pg, 500 pg, and 1 ng (n = 3 for each concentration) of MEHMP was placed at the bottom of a 1.5 mL vial. A 5-mm piece of PDMS tubing was exposed to the headspace for 18 h at 28 °C. Points represent sampling replicates; red: regression line. Based on the regression line, we estimate that a peak with an area of ca. 28.000 (arbitrary units; y) is equivalent to 200 pg of MEHMP (x).

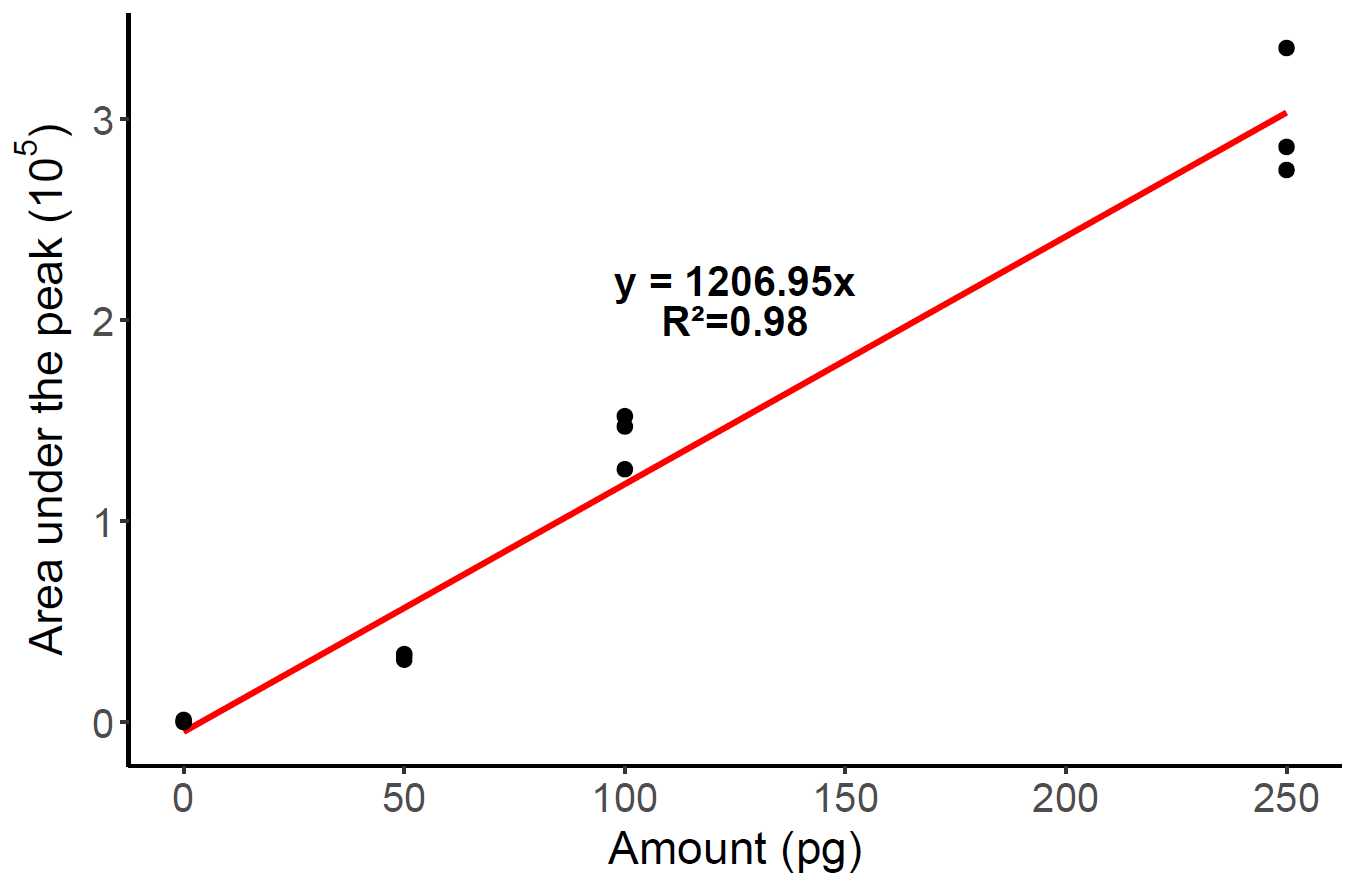

**Figure S3.** Calibration curve of methyl 3-ethyl-4-methyl-2-hydroxy-pentanoate (MEHMP) using SPME fiber and GC-MS. 1 µL of a hexane solution containing 0 pg, 50 pg, 100 pg or 250 pg (n = 3 for each concentration) of MEHMP was placed at the bottom of a 1.5 mL vial. SPME fibers (DVB/CAR/PDMS) were exposed to the headspace for 18 h at 28 °C. Points represent sampling replicates; red: regression line. Based on the regression line, we estimate that a peak with an area of ca. 240.000 (arbitrary units; y) is equivalent to 200 pg of MEHMP (x).

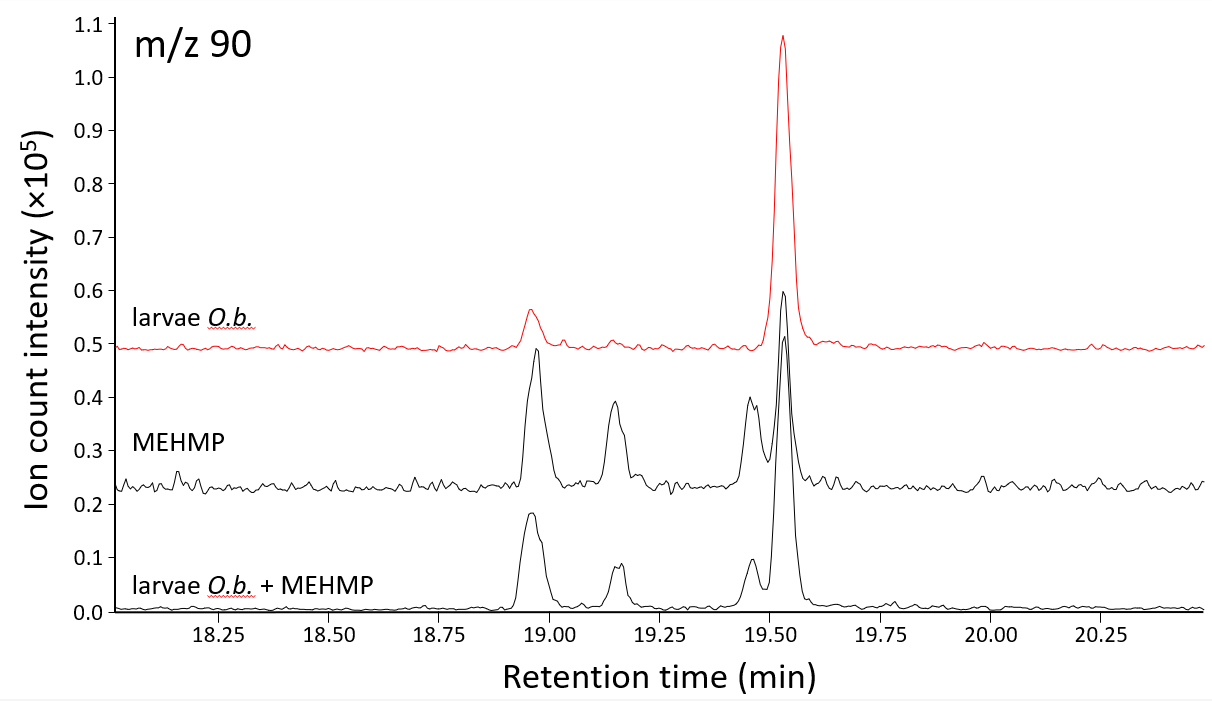
 **Figure S4.** GC-MS *m/z* 90 extracted ion chromatograms of SPME headspace extracts of *O. biroi* larvae (red, “larvae *O.b.*”), of the racemic mixture of synthesized MEHMP (“MEHMP”), and of *O. biroi* larvae headspace spiked with MEHMP (“larvae *O.b.* + MEHMP”). The synthetic MEHMP is composed of four stereoisomers, two of which are naturally produced by *O. biroi* larvae.

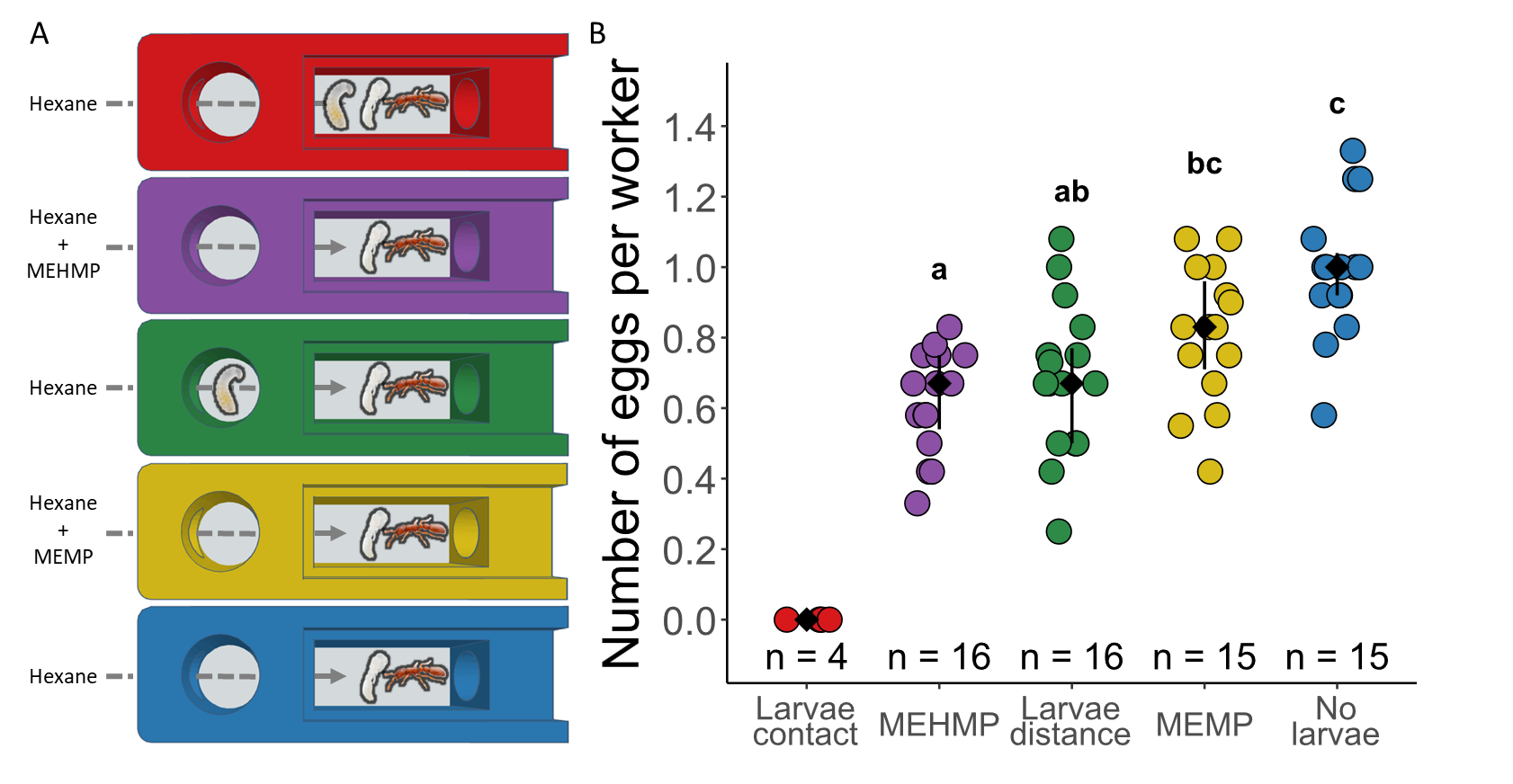

**Figure S5:** **Methyl 3-ethyl-2-hydroxy-4-methylpentanoate inhibits adult egg laying in a specific manner.** **(A)** 2-chamber arenas with controlled airflow. Treatments from top to bottom: workers in physical contact with larvae (red, “larvae contact”), workers exposed to synthetic MEHMP (purple, “MEHMP”), workers without physical contact with larvae (green, “larvae distance”), workers exposed to synthetic MEMP (yellow, “MEMP”), workers not exposed to larvae (blue, “no larvae”). Ants in all treatments were exposed to hexane and were in direct contact with pupae to ensure comparability among treatments. **(B)** Number of eggs per worker on the last experimental day. Data points represent replicate arenas. Black: median ± IQR. Different letters represent significantly different treatments (p < 0.05).
